# Constructing value on the fly from episodic memory supports flexible decision-making

**DOI:** 10.64898/2026.11.18.767634

**Authors:** Marco D’Alessandro, Giovanni Pezzulo

## Abstract

Value-based decisions often require people to flexibly draw on and combine information from past experiences to evaluate novel stimuli and adapt to changing demands. Although numerous studies have highlighted the involvement of episodic memory in value-based decision-making, whether and how a common episodic mechanism can support diverse decision-making scenarios remains poorly understood. Here, we introduce GENESIS-Value, a computational framework that constructs value on the fly by retrieving and integrating episodic memories, rather than relying on stable, precomputed value representations. The model combines generative stimulus representations with a flexible query–key–value retrieval mechanism, allowing the same memory store to support different valuation and choice strategies. Across five simulations, we demonstrate that GENESIS-Value can use episodic sampling to reconstruct the values of experienced stimuli, generalize to novel stimuli based on shared features, estimate feature values, guide both individual and sequential choices, and select among multiple offers involving different rewards and levels of risk. We further show that retrieval can be guided by stimulus features, temporal proximity, reminders, and reward outcomes, as well as combinations of these factors, enabling adaptation to different decision demands without requiring separate value functions. The model also captures qualitative behavioral signatures reported in empirical studies of episodic decision-making. Together, these findings establish a unified computational account of how episodic retrieval can support diverse forms of value-based inference and choice. Rather than replacing stable value representations, episodic retrieval may provide a complementary mechanism that allows decision-makers to flexibly address a variety of choice scenarios by reconstructing the value of available options on the fly.

## Introduction

How do we make value-based decisions? Classical neuroeconomic theories assume that we form relatively stable value representations of items or their attributes and compare these representations to make choices between offers, such as when choosing between foods or lotteries ^1,2^. The stable value can correspond, for example, to a running average of previous experiences, weighted by their recency ^3^. Similarly, reinforcement learning models assume that we select reward-maximizing actions, such as choosing between bandits or navigating the branches of a T-maze, based on learned value functions over states (or state– action pairs) ^4^. Candidate neural substrates for representing such stable economic values include cortical structures like the orbitofrontal cortex ^5–7^ and subcortical structures like the ventral striatum ^8–10^.

Stable value representations and value functions offer important computational advantages. They can be learned through trial and error, for example, when navigating an environment or choosing among bandits. Once learned, they provide a compact summary of past experience and allow the value of options to be computed in advance, reducing the computational demands of subsequent decisions. However, compressing experience into a fixed value function comes at the cost of flexibility. A value function learned to pursue a particular goal or maximize a particular type of reward may not generalize readily to novel situations in which the goal or choice context changes. For example, an agent might need to switch from selecting the option with the highest expected reward to selecting one that occasionally yields exceptionally high rewards or avoids punishment altogether, or prioritize some stimulus features over others in context-dependent ways. Accommodating such changes may require learning a new value function or maintaining multiple value functions in parallel. This leaves open the question of how humans maintain a high level of flexibility in value-based decision-making, adapting decisions to changing goals ^11–13^.

One possible solution is suggested by a growing body of research showing that episodic memory contributes to value-based decision-making. According to this account, decisions can be guided by retrieving and sampling from memory (even a small number of) past choice episodes, with the content of the retrieved episodes influencing the current choice ^14–29^. For example, previous studies showed that people retrieve detailed episodic memories to evaluate novel choices offers ^14^ and to guide decisions when the features relevant to future choices are uncertain ^17^, that human behavior during sequential choice (bandit) tasks can be well explained by an episodic sampling model, and that reminders of past choices bias sequential value-based decisions toward previously selected actions and their associated outcomes ^18^. Other studies suggest that episodic memory also plays a critical role in the Iowa Gambling Task (IGT) ^30^, a well-established experimental paradigm for studying decision-making under uncertainty, in which participants choose cards from multiple decks with different reward and penalty structures. Episodic memory may support these decisions by allowing people to flexibly update their estimates of deck values during the task ^31–34^. Furthermore, other studies have implicated the hippocampus, a structure strongly associated with episodic memory, in valuation and decision-making ^35–38^. Across multiple experimental paradigms, episodic retrieval may thus provide a means of flexibly evaluating current options in relation to specific past experiences, without requiring all relevant information to be integrated into single, fixed value representations.

Several theoretical frameworks have been proposed to explain how memory (including episodic memory) contributes to value-based decisions. The theory of dynamic memory and case-based reasoning ^39^ proposes that people draw on past experiences retrieved from episodic memory to form expectations and solve novel problems. Instance-based learning theory proposes that people solve decision-making tasks by learning and retrieving decision instances or examples ^40^. The theory of *decision by sampling* ^41^ proposes that people evaluate the value of an offer relative to a sample of other values retrieved from memory, such as the values of previously encountered offers. Related approaches incorporate episodic memories into reinforcement learning models, providing an alternative to both model-based planning and model-free choice. Such episodic mechanisms can be particularly effective early in learning, when conventional value functions have not yet been fully established ^42–44^. Other accounts similarly propose that value is reconstructed online from samples retrieved from memory at decision time, with retrieval selectively recruited when useful, such as under conditions of decision uncertainty, and the amount of retrieval effort adjusted to regulate the number of samples collected ^45–47^.

Despite these advances, at least three important questions remain. First, for episodic sampling to effectively support decision-making, it must provide accurate value estimates for both familiar and novel choice situations. However, the extent to which episodic value construction can achieve such accuracy, and how accuracy depends on computational resources, particularly the number of retrieved samples, remain poorly understood. Second, existing episodic models have largely been developed to account for specific tasks, and a comprehensive framework that captures the contribution of episodic memory across diverse decision-making scenarios, such as those examined in the experimental paradigms described above, is still lacking. Such a framework should incorporate a flexible retrieval mechanism that can select relevant episodes based on stimulus or feature similarity, temporal proximity, reward outcomes, or combinations of these factors. Third, existing approaches have largely been developed and evaluated using simplified stimuli and task settings, leaving open the question of how well episodic models generalize to more naturalistic stimuli, such as images. These stimuli have rich perceptual and semantic structures that provide multiple, complementary retrieval cues and more closely reflect the complexity of the information people encounter in everyday decision-making.

Here, we address these challenges by introducing GENESIS-Value (Generative Episodic–Semantic Integration System for Valuation), an episodic framework for memorizing and retrieving episodes associated with naturalistic stimuli, such as images. Rather than compressing past experiences into a fixed value function, GENESIS-Value constructs value on the fly by sampling episodic memories. Its flexible querying mechanism allows the same memory store to support diverse decision-making scenarios, while the number of samples determines how much episodic evidence is retrieved for each decision. The framework follows the principle of retrieval-augmented generation (RAG) ^48^, using retrieved memories to inform value construction. Crucially, episodes are stored as key–value associations, where each key serves as an index for retrieving a specific memory, much like a library catalogue entry identifies a book. Retrieval is guided by a query–key–value mechanism: a query (e.g., a request for a specific philosophy book, for philosophy books in general, or for Italian philosophy books of the fifteenth century) matches relevant keys, which in turn provide access to the associated memories. This mechanism flexibly selects episodes according to current decision requirements, allowing the same memory store to support different goals and choice strategies through semantic, temporal, or reward-based queries ^49,50^. For example, just as a library user can search for books with more or less specific features, an agent can prioritize recently rewarded options or retrieve episodes associated with gains rather than losses, without maintaining separate value functions for each strategy.

Across five simulations, we investigate the ability of GENESIS-Value to support value-based inference and decision-making through episodic sampling, and examine how its behavior relates qualitatively to established empirical findings in humans. Rather than matching empirical data exactly, our primary question is whether GENESIS-Value can support the wide range of strategies demonstrated in human experiments. In the first simulation, inspired by the experiment of ^14^, we test whether the model can accurately reconstruct the values of previously experienced stimuli and generalize to novel stimuli by inferring their values from features shared with previously encountered stimuli. In the second simulation, inspired by the experiment of ^17^, we examine whether the model can construct estimates of stimulus-feature values on the fly by sampling and combining episodic memories sharing these features, and use these estimates to guide adaptive choices. We also investigate how performance depends on the sampling budget. In the third and fourth simulations, inspired by the experiments of ^18^, we investigate whether the framework can support sequential decisions in two-armed bandit environments, and examine how reminders of past choices influence episodic retrieval and subsequent value-based decisions. Finally, in the fifth simulation, we turn to the Iowa Gambling Task (IGT) ^30^ to investigate whether GENESIS-Value can support adaptive choices among multiple offers involving different rewards and levels of risk.

## Results

### GENESIS-Value architecture

The GENESIS-Value architecture performs two main functions: *memorization* of choice episodes (here, Shape3D images and their associated rewards) and *retrieval* of these episodes to construct value representations on the fly before a choice (Figure 1). Following the GENESIS framework ^49^, these functions are implemented through two interacting components: a frozen conditional variational autoencoder (CVAE), which supports stimulus encoding and decoding, and a retrieval-augmented generation (RAG) mechanism, which supports the memorization and retrieval of episodic memories. These two interacting components can be putatively mapped to semantic and episodic processes, respectively.

**Figure 1:**
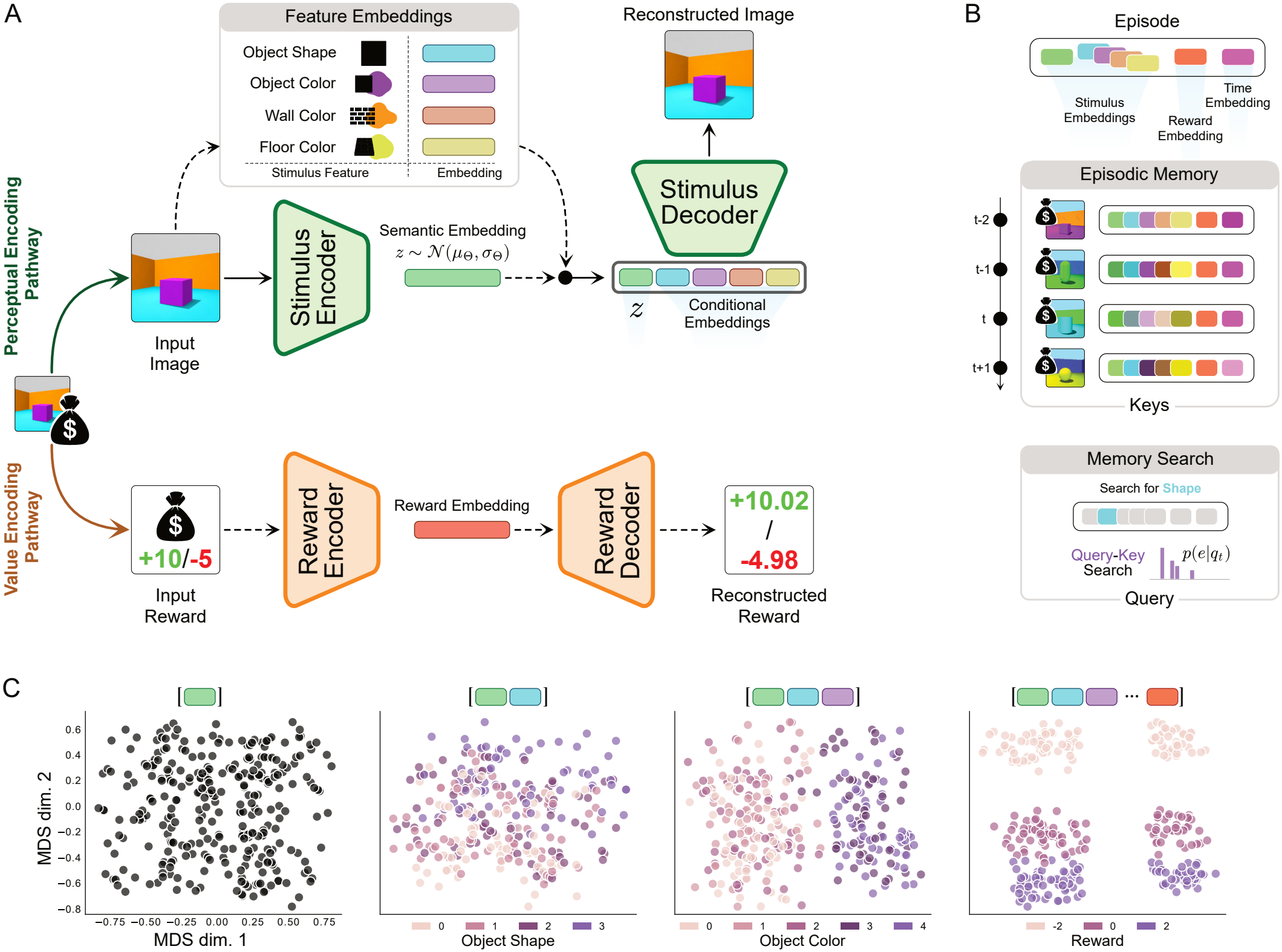
GENESIS-Value architecture. **A**. The GENESIS module processes each Shape3D image and its associated scalar reward along parallel stimulus- and reward-encoding pathways. A frozen CVAE encodes the image into a Gaussian semantic latent and conditional embeddings for object shape, object color, wall color, and floor color; the stimulus decoder combines these representations to reconstruct the original image. In parallel, the reward encoder maps the scalar reward to an embedding, from which the reward decoder reconstructs the input reward. **B**. The resulting stimulus and reward representations are combined with a temporal-context embedding to form an episode. Episodic memories are accessed through query–key matching. For visual clarity, only the searchable keys are depicted; each key provides access to an associated value containing the stored episodic content. **C**. Content of episodic memories projected in two dimensions through multidimensional scaling (MDS). The plots help appreciate the fact that episode composition reshapes the geometry of the memory representation. Enriching the episode by progressively including semantic-only (leftmost panel), perceptual (second and third panels), and reward (rightmost panel) embeddings entails a re-organization of the latent geometry.

During *memorization*, the frozen CVAE encodes each Shape3D image into an image latent *z* and learned embeddings representing its features, including object shape, object color, floor color, and wall color. The CVAE was pretrained using four object-shape classes (cube, sphere, cylinder, and pill) and five classes for each of the three color features (object color, floor color, and wall color). Unless otherwise specified, this pretrained CVAE was used across all Shapes3D-based simulations. A separately trained reward encoder creates an embedding of the reward associated with the same image. Each episode binds together these stimulus representations, the reward embedding, and a gradually evolving temporal-context embedding that indicates when the episode occurred (Figure 1A). More specifically, in GENESIS-Value, each episode is represented as a key–value association. The value contains the compressed stimulus and reward representations, while the key additionally incorporates a temporal-context embedding, providing contextual information independent of stimulus identity. This allows memories to be accessed through both stimulus-related and stimulus-independent information, while their associated values provide the content used for reconstruction and decision-making. For simplicity, the figures depict only keys, leaving their associated values implicit.

During *retrieval*, relevant episodes are recollected using the RAG query–key–value mechanism (Figure 1B). Note that, in this context, *value* refers to the content of a retrieved episode, not its economic value. The identity and number of the retrieved episodes depend on several factors, including the retrieval query, which can prioritize particular stimulus features, rewards, and/or temporal proximity, and the number of samples retrieved, as explored in the following simulations. The stimuli and rewards associated with the retrieved episodes are then decoded using the CVAE and reward decoder, respectively, and the retrieved rewards are aggregated to guide decision-making. For example, when choosing among multiple bandits, the model retrieves episodes associated with previous interactions with each bandit, aggregates their rewards, and compares the resulting estimates to identify the bandit with the highest estimated value.

Figure 1C schematically illustrates the content of episodic memories formed from images varying in shape, color, and reward, projected in two dimensions through multidimensional scaling (MDS) (other types of content, such as temporal information, are omitted for simplicity). The panels show how the latent geometry of episodic memory – namely, the relative distances between stimulus representations in a low-dimensional space, reflecting their similarity in terms of encoded features – changes as different types of information are incorporated, from semantic features alone (leftmost panel) to perceptual features (second and third panels) and reward information (rightmost panel). More broadly, Figure 1C helps appreciating the fact that episodic memories occupy a high-dimensional space, in which each dimension represents a feature of experience (e.g., stimulus identity, object shape and color, floor color, wall color, or reward) and episode composition reshapes the geometry of the memory representation. The similarity between episodes depends on which features are emphasized by the retrieval query: for example, a query emphasizing color makes episodes with similar colors more relevant, whereas a reward-based query prioritizes episodes with similar reward outcomes. In this sense, each query defines a task-relevant projection of the memory space by emphasizing selected features while down-weighting others. Thus, the same memory store can potentially support distinct value-based decision strategies through flexible retrieval, without requiring separate value representations for each strategy.

### Episodic sampling accurately reconstructs the value of experienced stimuli and generalizes to novel stimuli

As a first step in the evaluation of episodic retrieval in GENESIS-Value, we asked whether it can accurately reconstruct the value associated with seen stimuli and generalize to infer the value of novel, unseen stimuli that share features with seen stimuli. To investigate this, we adapted the task developed by Barron et al. ^14^ , in which participants chose between novel foods created by combining familiar ones. The study showed that participants constructed appropriate values of these novel stimuli by integrating multiple independent memories, with the hippocampus and medial prefrontal cortex implicated in this process.

We constructed 125 Shape3D cube images by independently varying three features: object color, floor color, and wall color, across five levels each (5^3^ possible conjunctions). Each color level was assigned a contribution in ™2, ™1, 0, 1, 2, and the ground-truth value of a stimulus was defined as the mean contribution of its three color features, *V*^⋆^(*s*) = (*v*_object_ + *v*_floor_ + *v*_wall_)/3.

In each of 125 independently seeded sessions, one image was selected for every conjunction and one of them was designated as the target stimulus. We evaluated two conditions using the same target stimulus: a *seen* condition, in which the target and its reward were included among all 125 stored episodes, and an *unseen* condition, in which the target was withheld, leaving 124 reward-bearing episodes in memory (Figure 2A). Thus, the two conditions differed only in whether the complete target conjunction and its associated reward were available in episodic memory. In both conditions, the target stimulus was used to form a retrieval query combining its image latent and its object-, floor-, and wall-color representations with equal weight, with sampling temperature fixed at *τ* = 0.25. The target reward was never included in the query.

**Figure 2:**
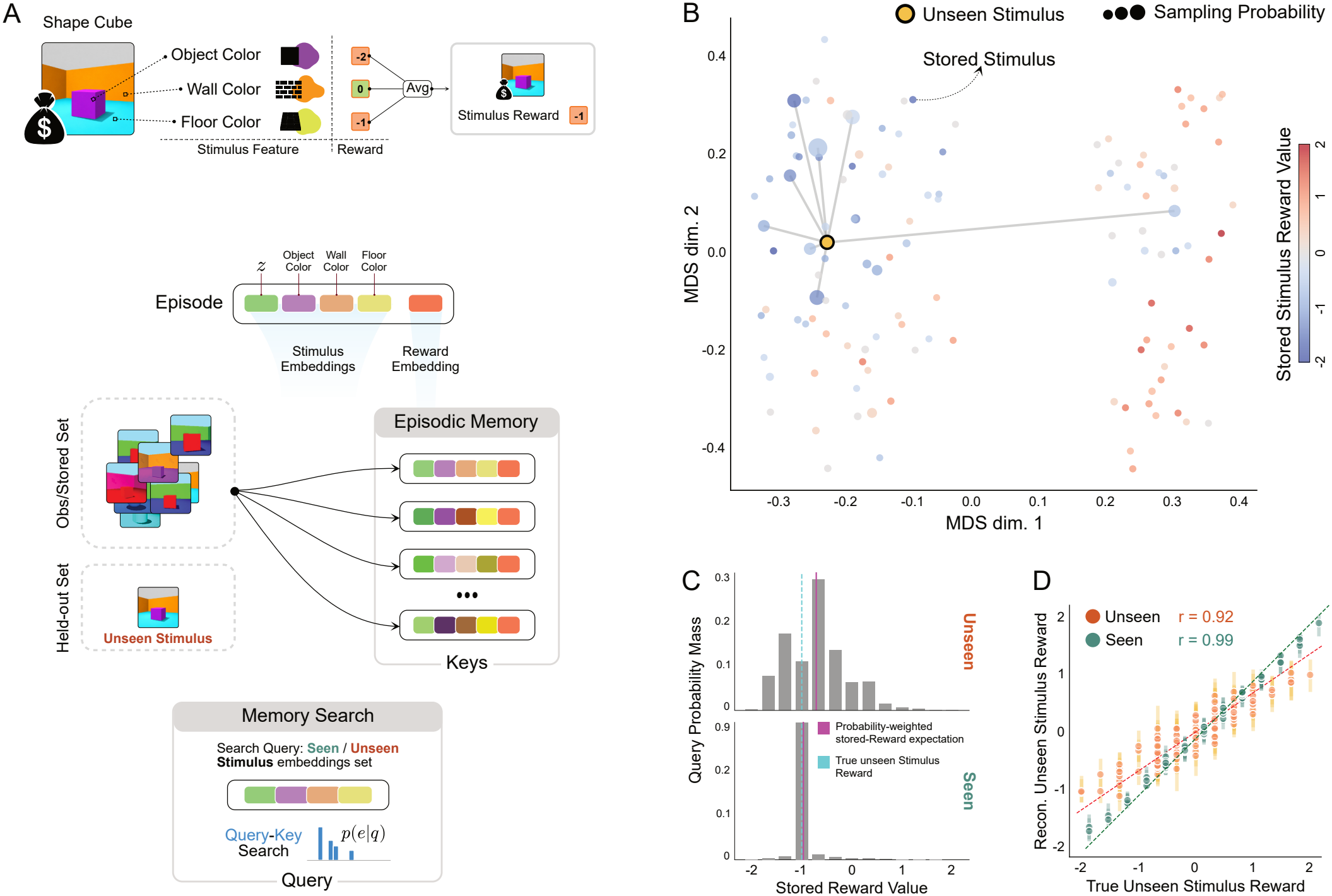
Episodic sampling reconstructs experienced stimulus values and generalizes to novel stimulus conjunctions. **A**. Schematic of stimulus-value reconstruction in GENESIS-Value, illustrated for a novel stimulus. A stimulus defined by a previously unseen (and never stored) conjunction of object, floor, and wall colors provides a perceptual-semantic retrieval query. The query assigns accessibility to stored reward-bearing episodes according to stimulus similarity, and sampled reward embeddings are aggregated to construct the stimulus value. **B**. Query-dependent episodic geometry for an example held-out stimulus. The 124 stored conjunctions and the novel target are projected into two dimensions from their weighted stimulus representations using classical multidimensional scaling. Stored-point color indicates experienced reward and point size indicates the query-specific sampling probability computed by GENESIS-Value. Lines connect the target to the eight episodes with the highest sampling probabilities. For this target, retrieval concentrated on feature-overlapping, negatively valued memories. **C**. Query-weighted reward evidence for the same target in the unseen (top) and seen (bottom) conditions. Bars show the total sampling probability assigned to stored episodes at each reward value. In the unseen condition, retrieval probability was concentrated on negative-valued memories, yielding a probability-weighted expectation of 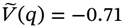. In the seen condition, the exact target episode received the highest sampling probability (*p* = 0.89), shifting the expectation to 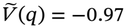, close to the true value of ™1. These quantities provide scalar summaries of the reward evidence implied by the retrieval distributions; the GENESIS-Value reconstruction itself operates by sampling reward embeddings, averaging them, and decoding the resulting aggregate. **D**. Value reconstruction across 125 independently seeded sessions at sampling budget *k* = 20, comparing seen (green) and unseen (orange) stimuli. Each point plots the true stimulus value against the mean value reconstructed from 500 stochastic retrieval repetitions; vertical intervals denote the 2.5th–97.5th percentiles across repetitions, and the dashed diagonal denotes perfect reconstruction. Fitted lines show the relationship between true and reconstructed values for each condition. Reconstruction was strongly correlated with true value for seen stimuli (*r* = 0.99) and remained strongly correlated for unseen stimuli (*r* = 0.92), although the latter exhibited greater compression toward the center of the reward distribution.

Figure 2B illustrates the geometry of episodic memory through a two-dimensional projection, obtained using multidimensional scaling (MDS), of the distances between stored episode keys. Each dot represents a stored episode, with color indicating its associated reward and size indicating its sampling probability given the target query, computed from the full weighted query–key similarities. The figure also shows an example unseen target (in yellow) with *V*^⋆^ = −1. The unseen target occupied a region of the query-relevant stimulus space close to a subset of stored episodes, which consequently received larger sampling probabilities. The position of each stimulus in this space depends on its particular combination of features, such that stimuli sharing more features tend to be located closer together. In this example, the unseen target lies near stored episodes sharing some of its features, which are predominantly associated with negative rewards. Although the exact target stimulus is not stored in episodic memory and therefore cannot be directly retrieved, GENESIS-Value can sample similar episodes and use their associated rewards to infer the value of the unseen target.

We examined whether query–based episodic retrieval allowed reconstructing the value of a target stimulus with *V*^⋆^ = −1, both when it was seen and unseen (Figure 2C). To illustrate how the retrieval process provides evidence for value reconstruction, we consider a representative example from a single simulation instance. We examined the distribution of sampling probabilities across the reward values associated with stored episodes, separately for the seen and unseen conditions. For visualization, the probabilities of memories carrying the same reward across all sessions were summed to obtain 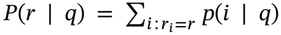, and the corresponding probability-weighted stored-reward expectation was

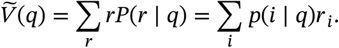

For the example target, when the stimulus was seen, its exact episode received the highest sampling probability, and the expected value approached the true reward 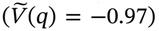. Thus, the availability of an exact episodic precedent leads to an accurate reward estimation. When the same stimulus was novel, retrieval probability was concentrated on episodes with negative values, yielding 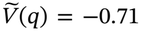. This estimate had the same sign as the true target value, but was closer to zero. This “regression to the mean” of the distribution of stimulus values (which was zero) follows from the similarity-based nature of the reconstruction: because the exact conjunction is absent from memory, its value must be inferred from partially overlapping episodes whose rewards are generally less extreme than that of the unseen target.

We then tested whether these relationships generalized across 125 independently seeded sessions, with stimulus values ranging from −2 to 2. For each session and condition, episodes were sampled with replacement from the query-specific distribution, their reward embeddings were averaged and decoded, and reconstruction was repeated 500 times. Figure 2D shows the resulting stimulus-value estimates at *k* = 20, plotting the mean reconstruction against the true target value for both seen and unseen stimuli. When the target was seen, reconstructed values closely matched their true values (*r* = 0.99), and when the target was unseen, reconstructed and true values remained strongly correlated (*r* = 0.92), indicating that queries associated with more positively valued targets systematically retrieved more positive reward evidence, whereas negatively valued targets produced more negative estimates. Extreme positive and negative values were reconstructed closer to zero, showing the “regression to the mean” discussed above.

Taken together, these results show that GENESIS-Value supports both accurate value reconstruction for experienced stimuli and generalization to novel stimuli with similar features, qualitatively reproducing the findings of Barron et al. ^14^ . These findings support the idea that stimulus values can be reconstructed accurately through episodic sampling alone.

### Episodic sampling supports feature-conditioned value estimation and choice

We next asked whether GENESIS-Value can estimate the value associated with a particular feature by sampling and integrating episodic memories that share that feature, and whether it can use this information to guide novel decisions. To address this question, we adapted the task developed by Nicholas and Mattar ^17^ , in which participants first encoded stimuli composed of multiple features (e.g., color and category) and were later presented with offers defined by a single feature, which they had to accept or decline (e.g., accepting or declining all red items). The study suggests that when participants could not anticipate which feature would be relevant for future choices, they memorized the stimuli in full detail, retaining information about all their features. At decision time, they then retrieved the relevant past experiences (i.e., those involving stimuli that shared the target feature) and combined the rewards associated with those experiences.

We first tested whether GENESIS-Value supports feature-conditioned value estimation in 20 independently seeded sessions, each containing 200 Shape3D images varying along four features: shape (the main object of our analysis), object color, floor color, and wall color. Each stimulus could show one of four object shapes (e.g. cube, sphere, cylinder, capsule). These shapes occurred equally often, while object color, floor color, and wall color were balanced within each shape. Each episode was assigned one of four rewards, −2, −1, +1, or +2, with equal frequency across the session. Reward assignments preserved a session-wide mean of zero, but episodes containing a cube had a positive mean reward of 0.55 (with a range of 0.50–0.64 across sessions).

Each stimulus was stored in memory as an episode containing all its features and the associated reward (Figure 3A). At retrieval, the *cube* object shape was used as the query (Figure 3B), producing a strongly non-uniform accessibility distribution over the complete episodic memory (Figure 3E). Sampling temperature was fixed at *τ* = 0.25, and the sampling budget was varied over

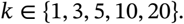

**Figure 3:**
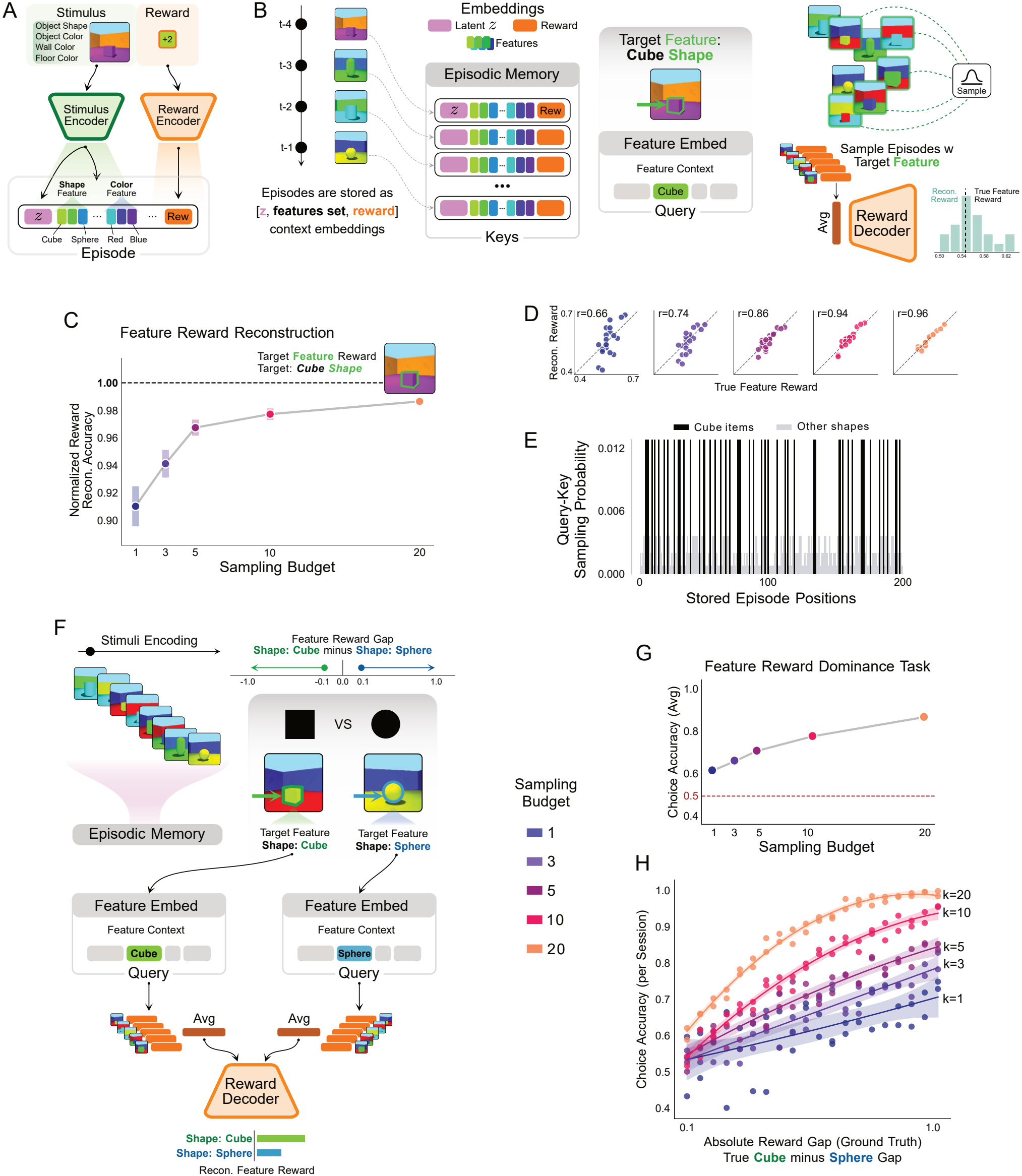
Feature-conditioned episodic sampling supports feature value reconstruction and flexible choice. **A**. Feature-rich Shape3D events are stored as episodes containing stimulus information and the associated reward. **B**. A task-relevant semantic feature selectively cues episodic sampling, and the rewards carried by retrieved episodes are combined to reconstruct a feature-conditioned value. **C**. Ground-truth-normalized cube-reward reconstruction accuracy across sampling budgets. Accuracy is the across-session mean of 100 × (1 − |reconstruction error|/realized cube reward); error bars show bootstrap standard errors across sessions. **D**. Session-level reconstructed versus true cube value across sampling budgets, showing increasing concentration around the identity relation as sampling budget increases. **E**. Example retrieval distribution induced by a cube query, showing highly selective but non-exclusive access to cube episodes. **F**. Flexible feature comparison, motivated by Nicholas and Mattar ^17^ , in which independently reconstructed cube and sphere values are compared to guide choice. **G**. Mean feature-choice accuracy as a function of sampling budget; the dashed line denotes chance. **H**. Session-level choice accuracy as a function of the absolute true cube–sphere value difference, showing greater reliability for easier comparisons and larger sampling budgets.

Sampling was performed with replacement and each session-by-budget condition was repeated 500 times. Under these conditions, at least 99.1% of sampled episodes contained the queried cube feature at every sampling budget, supporting the validity of the episodic sampling process.

GENESIS-Value estimated the value associated with the queried feature (e.g., shape cube) as the average reward across sampled episodes, with reliability increasing as the number of retrieval draws increased. Ground-truth-normalized reconstruction accuracy improved with *k* (Figure 3C), reflecting a monotonic reduction in session-level reconstruction error. The same effect was apparent at the level of individual sessions: estimates obtained from few retrieval draws were relatively dispersed around their corresponding ground-truth values, whereas estimates became progressively concentrated around the identity relation as more samples were averaged (Figure 3D).

Having established feature-conditioned value estimation, we next asked whether the same mechanism could support an accurate choice between two features. Taking inspiration by the task developed by Nicholas and Mattar ^17^ , we assumed that decision-makers would first encode feature-rich episodes in memory and then asked to make decisions based on one of the features. Specifically, here the task was to determine which of two object shapes, *cube* or *sphere*, had the larger value in the current episodic history (Figure 3F).

We generated 40 additional 200-event sessions, comprising 20 cube-dominant and 20 sphere-dominant sessions. Decision difficulty was controlled by the signed feature-reward difference

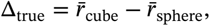

whose absolute magnitude was evenly spaced from 0.1 to 1.0 within each dominance condition. On every simulated decision, independent cube and sphere queries produced two reconstructed values, and the feature with the larger value was selected. The same sampling budgets were evaluated at *τ* = 0.25, with replacement sampling and 500 paired decisions per session and budget.

Choice accuracy remained above chance across the configured sampling budgets and increasing the number of samples substantially improved the probability of selecting the feature with the larger true value (Figure 3G). Performance also depended on decision difficulty: sessions with larger absolute cube–sphere value differences were classified more accurately, while larger *k* improved choice accuracy across the controlled range of value gaps (Figure 3H).

In summary, these simulations show that GENESIS-Value can support feature-based decisions with episodic sampling, as demonstrated empirically in the study of Nicholas and Mattar ^17^ . GENESIS-Value can already support these decisions with just one sample, but its performance improves as the number of samples increases. This result shows that value estimation can be accurately realized through episodic sampling – by retrieving relevant past episodes based on the features that matter for the current decision and inferring value from their associated outcomes – rather than by maintaining a separately learned value for every possible stimulus feature.

### Episodic sampling supports sequential choices

We next asked whether GENESIS-Value can support sequential decision-making. To assess this capability, we adapted the task developed by Bornstein et al. ^18^ , in which participants performed a sequential multi-armed bandit task, with reward probabilities changing over time. The study showed that an episodic model better explained human choices than a standard reinforcement-learning (RL) model based on a recency-weighted (through learning rate) average of past rewards. Here, our research questions were different: rather than comparing the ability of episodic and RL models to explain human choices, we asked whether GENESIS-Value can achieve performance comparable to temporal-difference (TD) learning in volatile two-armed bandit tasks by using temporal context to guide retrieval, and whether adjusting the sampling temperature, which controls the temporal range of retrieved episodes, allows the model to adapt to different levels of environmental volatility.

We evaluated GENESIS-Value in two-armed bandits in which each experienced trial was stored as an episode containing the selected action, a binary outcome *r*_*t*_ ∈ {−1, +1}, a trial-unique Shape3D stimulus representation, and temporal context. At each subsequent choice, the model reconstructed one value for each arm from arm-specific episodic samples and converted their difference into a stochastic choice.

The sampling budget was fixed at *k* = 20, while sampling temperature *τ*, which controls retrieval selectivity, was varied across

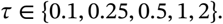

By design, the temperature manipulation changes the effective temporal horizon of the evidence used for valuation: lower temperatures concentrate sampling on the episodes whose temporal contexts most closely matched the present state, whereas higher temperatures flatten the distribution and allow more distant episodes to contribute.

We simulated 100 independently generated reward environments with 50 agents in each environment. Every environment comprised 130 choice trials. The probability of a positive reward (*r*_*t*_ = +1) for the left arm followed a reflected Gaussian random walk bounded between 0.3 and 0.7, while the right-arm probability was its complement. Agents selected randomly during the first 20 trials; thereafter, temporal context was the sole retrieval query for reconstructing the two action values (Figure 4A,B)

**Figure 4:**
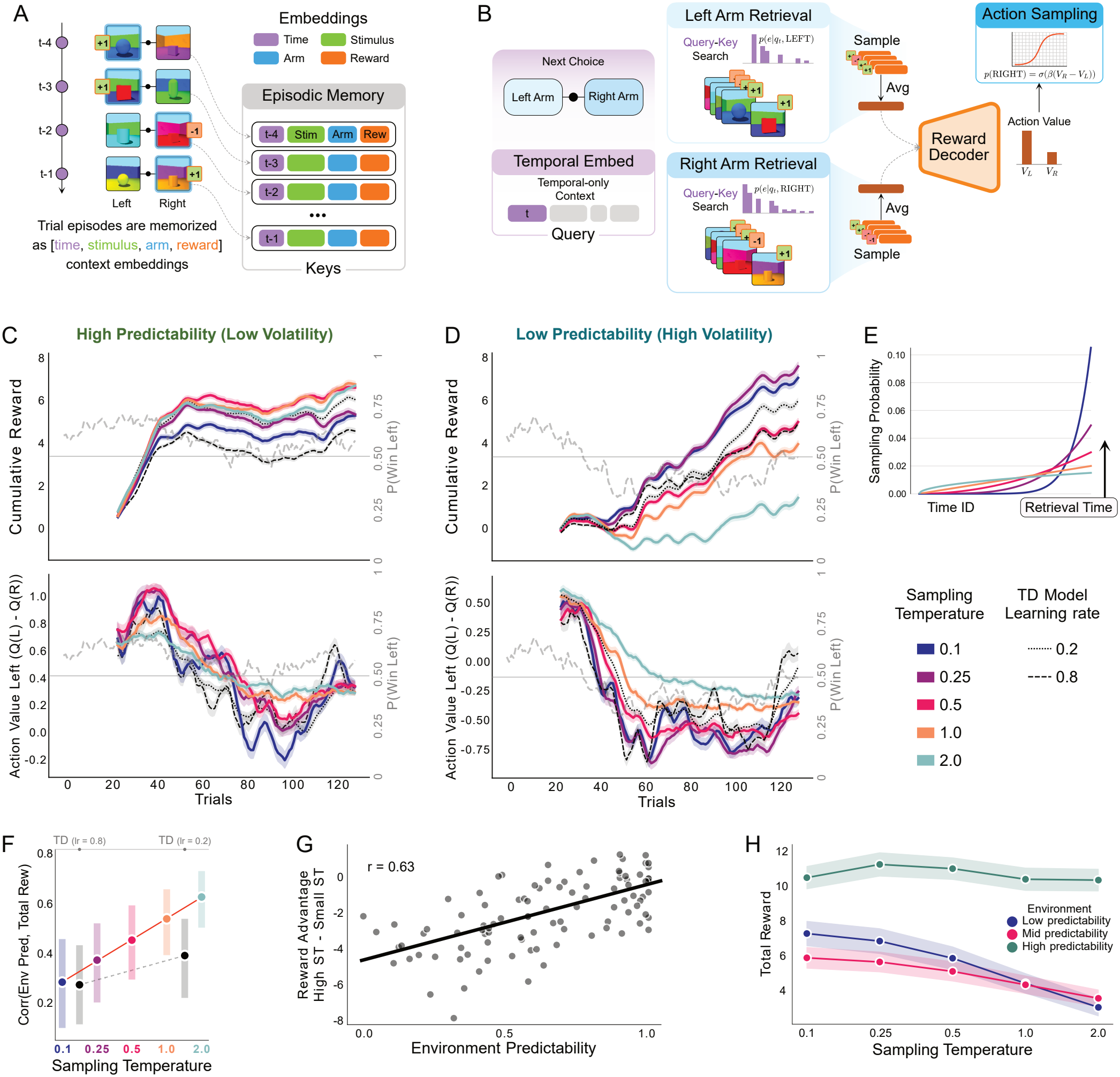
Sampling temperature determines how environmental predictability shapes reward-guided performance. **A**. Two-armed bandit task. Each choice yields a binary outcome and a trial-unique Shape3D stimulus; action, reward, stimulus, and temporal context are stored as one episode. **B**. Episodic valuation. Current temporal context queries arm-specific memories. Twenty episodes are sampled with replacement for each reconstructed action value, while sampling temperature *τ* controls retrieval selectivity. **C**. Representative high-predictability, low-volatility environment. Traces show cumulative reward and reconstructed left-minus-right action values across GENESIS-Value sampling temperatures and for temporal-difference models with learning rates *α* = 0.8 and *α* = 0.2; the gray dashed trace is the left-arm reward probability. High-temperature sampling benefits from distributed evidence that remains informative. **D**. Representative low-predictability, high-volatility environment. The panel also includes temporal-difference models with learning rates *α* = 0.8 and *α* = 0.2. The pattern reverses:low-temperature, recency-selective sampling better tracks changing contingencies, whereas high-temperature sampling incorporates increasingly obsolete outcomes. **E**. Example query–key sampling distributions across temperatures. Low-temperature sampling preserves a strong recency gradient, whereas increasing temperature flattens the distribution and increases access to temporally distant episodes. **F**. Across-environment correlation between predictability and total reward for the GENESIS sampling-temperature conditions and temporal-difference models with learning rates *α* = 0.8 and *α* = 0.2. Increasing temperature strengthens the dependence of performance on persistent historical evidence. **G**. Reward advantage of high-temperature over low-temperature sampling, defined as mean(*τ* = 1, 2) − mean(*τ* = 0.1, 0.25), as a function of environmental predictability. Negative values favor low-temperature sampling; positive values favor high-temperature sampling. **H**. total reward across sampling temperatures for low-, intermediate-, and high-predictability environments. Low-temperature sampling is favored at lower predictability, whereas high-temperature sampling becomes increasingly beneficial as reward structure persists. Lines and shaded regions show means and uncertainty across environments.

At *τ* = 0.1 (low temperature), episodic sampling in GENESIS-Value was strongly concentrated on episodes temporally closer to the present, producing a pronounced recency bias. Increasing *τ* toward higher sampling temperatures progressively flattened this gradient and increased access to temporally distant episodes (Figure 4E). We therefore asked when broad temporal integration was useful by assigning each reward environment a predictability score that quantified how consistently the direction of the early reward advantage identified the better arm later in the same environment. High scores indicate that older evidence remains informative; low scores indicate that the identity of the better arm changes more frequently, therefore reflecting that early reward advantages are less informative about which arm is better later.

Representative environments revealed opposite effects of sampling temperature depending on reward dynamics (Figure 4C,D). In a highly predictable environment, high-temperature sampling integrated temporally distributed but mutually consistent evidence, increasing cumulative reward relative to low-temperature sampling (Figure 4C). In a weakly predictable environment, older outcomes were more often obsolete, and low-temperature sampling better tracked the locally current reward structure (Figure 4D; cumulative rewards should be compared across temperatures within, rather than between, environments, given their different reward dynamics).

The interaction between sampling temperature and environmental predictability in GENESIS-Value generalized across the 100 environments considered in our analysis. The correlation between predictability and total reward increased with sampling temperature (Figure 4F), indicating that performance under high-temperature sampling depended more strongly on whether temporally distant evidence remained informative.

As a non-episodic control, we also implemented temporal-difference (TD) learning ^51^, a standard model-free reinforcement-learning method also used as a reference model by Bornstein et al. ^18^ . We considered TD learners with high (*α* = 0.8) and low (*α* = 0.2) learning rates. Because a larger learning rate places greater weight on recent outcomes, the *α* = 0.8 learner has a shorter effective integration timescale than the *α* = 0.2 learner. Across environments, the correlation between predictability and total reward was lower for *α* = 0.8 than for *α* = 0.2 (Figure 4F). This ordering parallels the GENESIS-Value results: increasing the sampling temperature in GENESIS-Value and decreasing the learning rate in TD both broaden the temporal span over which reward information influences choice, making performance more dependent on the relevance of past outcomes. Both TD correlations nevertheless remained below those observed for the higher-temperature GENESIS-Value conditions (see also example individual trajectories for the two TD learners in Figures 4C,D).

Finally, we summarized the interaction between sampling temperature and environment predictability in GENESIS-Value with the contrast

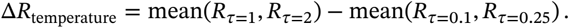

The contrast changed sign across the predictability range (Figure 4G): low-temperature sampling was advantageous when early reward evidence poorly predicted later contingencies, whereas high-temperature sampling became advantageous as reward direction became more persistent. Grouping environments into low-, intermediate-, and high-predictability classes yielded the same interaction (Figure 4H). Sampling temperature therefore regulates the effective temporal scale over which episodic reward evidence is integrated.

In summary, this simulation shows that GENESIS-Value can support sequential decision-making in bandit tasks, consistent with the findings of Bornstein et al. ^18^ . By adjusting the sampling temperature at decision time, the model can adapt to different levels of environmental volatility, producing a range of behavioral outcomes comparable to those achieved by TD learning with different learning rates. However, whereas TD learning requires setting the learning rate parameter during learning, GENESIS-Value can adjust its sampling temperature at decision time, allowing it to respond more readily to changing environmental demands.

### Episodic sampling reconstructs value under the influence of reminder cues

We next asked whether GENESIS-Value can flexibly use its query–key matching mechanism to guide episodic retrieval and value construction toward specific experienced stimuli. To investigate this, we adapted the task developed by Bornstein et al. ^18^ , in which participants performed the sequential bandit task described in Section , with explicit reminders of previously encountered stimuli provided on some trials. The study showed that these reminders can bias subsequent choices toward the reminded stimuli and their associated values.

We modified the bandit task of Section , in such a way that selected choice trials were preceded by a reminder stimulus: a held-out Shape3D image from the same complete semantic class as the stimulus associated with an earlier target trial, but perceptually distinct from the originally experienced image (Figure 5A). Using a held-out exemplar required retrieval based on learned semantic representations rather than visual identity, thereby making the process more challenging.

**Figure 5:**
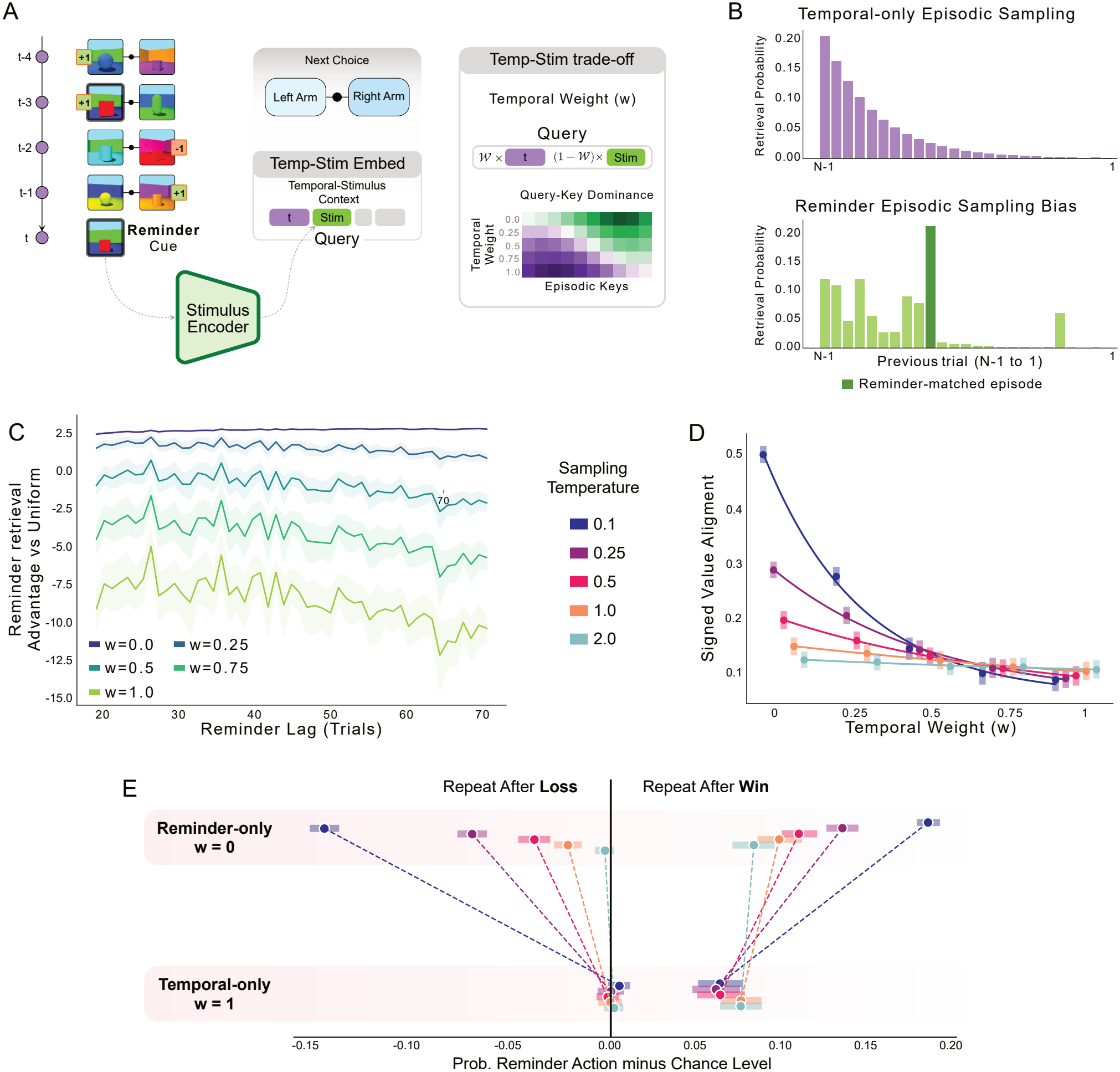
Reminder cues redirect episodic valuation. **A**. Reminder-bandit task. Before selected choices, the agent receives a novel image from the same Shape3D semantic class as the stimulus associated with an earlier target episode. The reminder is combined with current temporal context to form a retrieval query, with temporal weight *w* controlling their relative contribution. **B**. Example accessibility distributions. A temporal query preferentially accesses recent episodes, whereas reminder information reallocates probability toward the semantically matched target. Retrieval uses 20 samples with replacement. **C**. Target retrieval advantage relative to uniform sampling as a function of reminder lag, defined as the log ratio between target sampling probability and its uniform probability among eligible candidates. Reminder-dominated retrieval preserves access to distant targets, whereas increasing temporal weight strengthens lag dependence. **D**. Signed alignment between the reconstructed action-value difference and the target episode’s action–outcome content across temporal weights and sampling temperatures. Alignment declines strongly with temporal weight under low-temperature sampling, but this dependence becomes progressively shallower as temperature rises; at *τ* = 2, alignment is nearly invariant to query composition. **E**. Change in target-action repetition probability relative to 0.5. Left: following target losses, reminder-only retrieval promotes switching, most strongly at low sampling temperatures, whereas temporal-only retrieval approaches chance. Right: following target wins, reminder-only retrieval promotes repetition, again most strongly under low-temperature sampling; temporal-only retrieval reduces but does not eliminate this bias. Colors denote sampling temperature, horizontal intervals show 95% bootstrap intervals across independent environments, and red hatching marks intervals crossing the no-preference reference.

We then tested whether and to what extent incorporating the reminder into the retrieval query would bias episodic sampling, value reconstruction, and subsequent choices. We used *k* = 20 replacement sampling and the temperature range used above

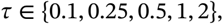

and manipulated the composition of the retrieval query. The query combined reminder-stimulus similarity with current temporal-context similarity according to a temporal weight

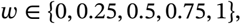

where *w* = 0 produced reminder-only retrieval and *w* = 1 produced the temporal-only retrieval. Reminder targets occurred 20– 70 choice trials in the past. Fifty independent environments with 50 agents per environment were simulated. Thus, *w* controlled which information defined episodic relevance, whereas *τ* controlled how selectively those relevance differences were translated into retrieved samples.

A reminder substantially redistributed retrieval probability toward its target episode relative to a temporal-only query (Figure 5B). Target accessibility was quantified relative to the uniform probability among eligible memories. When the reminder dominated the query, the target retained a strong accessibility advantage despite temporal distance. Increasing temporal weight restored the influence of recency, producing a stronger decline of target accessibility with reminder lag; under temporal-only retrieval, sufficiently distant target episodes fell below the uniform reference (Figure 5C).

We next examined how this accessibility change propagated to reconstructed value. The right-minus-left value difference was sign-oriented by the target action and target outcome, so positive alignment indicated a value configuration consistent with repeating the target action after a win or avoiding it after a loss. At low sampling temperatures, value alignment declined strongly as temporal weight increased: a reminder-dominated query gave the target episode substantial leverage over reconstructed value, but this leverage rapidly weakened as temporal context took over the query. The dependence on query composition became progressively shallower as *τ* increased and the retrieval distribution flattened (Figure 5D).

Finally, these reconstructed values altered the immediately subsequent choice (Figure 5E). Following target losses, reminderdominated retrieval promoted switching away from the reminded action, with the strongest effect under low-temperature sampling; temporal-only retrieval approached chance-level repetition. Following target wins, reminder-dominated retrieval increased repetition of the target action, again most strongly at low temperatures. The reminder-to-temporal difference therefore decreased as the sampling temperature increased.

In summary, this simulation shows that incorporating reminders of past episodes into the retrieval query biases episodic retrieval, value construction, and subsequent choices toward the reminded episode, qualitatively reproducing the empirical findings of Bornstein et al. ^18^ . Comparing temporal-only and reminder-based sampling illustrates how the query–key matching mechanism in GENESIS-Value can flexibly guide episodic retrieval and value construction using the same underlying memories.

### Episodic sampling supports adaptive choices in the Iowa Gambling Task

Finally, we tested whether GENESIS-Value can support adaptive choices among multiple offers involving different rewards and levels of risk, and whether different retrieval queries – prioritizing, for example, temporal information, sensitivity to rewards and losses, or combinations of these factors – can give rise to different choice strategies. To address this question, we adopted the Iowa Gambling Task (IGT): a standardized neuropsychological assessment of decision-making under uncertainty, risk-taking, and sensitivity to rewards and punishments. In the IGT, participants repeatedly select cards from four decks that differ in both long-run return and the frequency and magnitude of punishments (Figure 6A) ^30^. Decks A and B provide larger immediate rewards but are disadvantageous over repeated choices, whereas Decks C and D provide smaller rewards but favorable long-run returns. The task also dissociates punishment frequency from overall value: Decks A and C contain relatively frequent punishments, while Decks B and D contain rarer but larger punishments.

**Figure 6:**
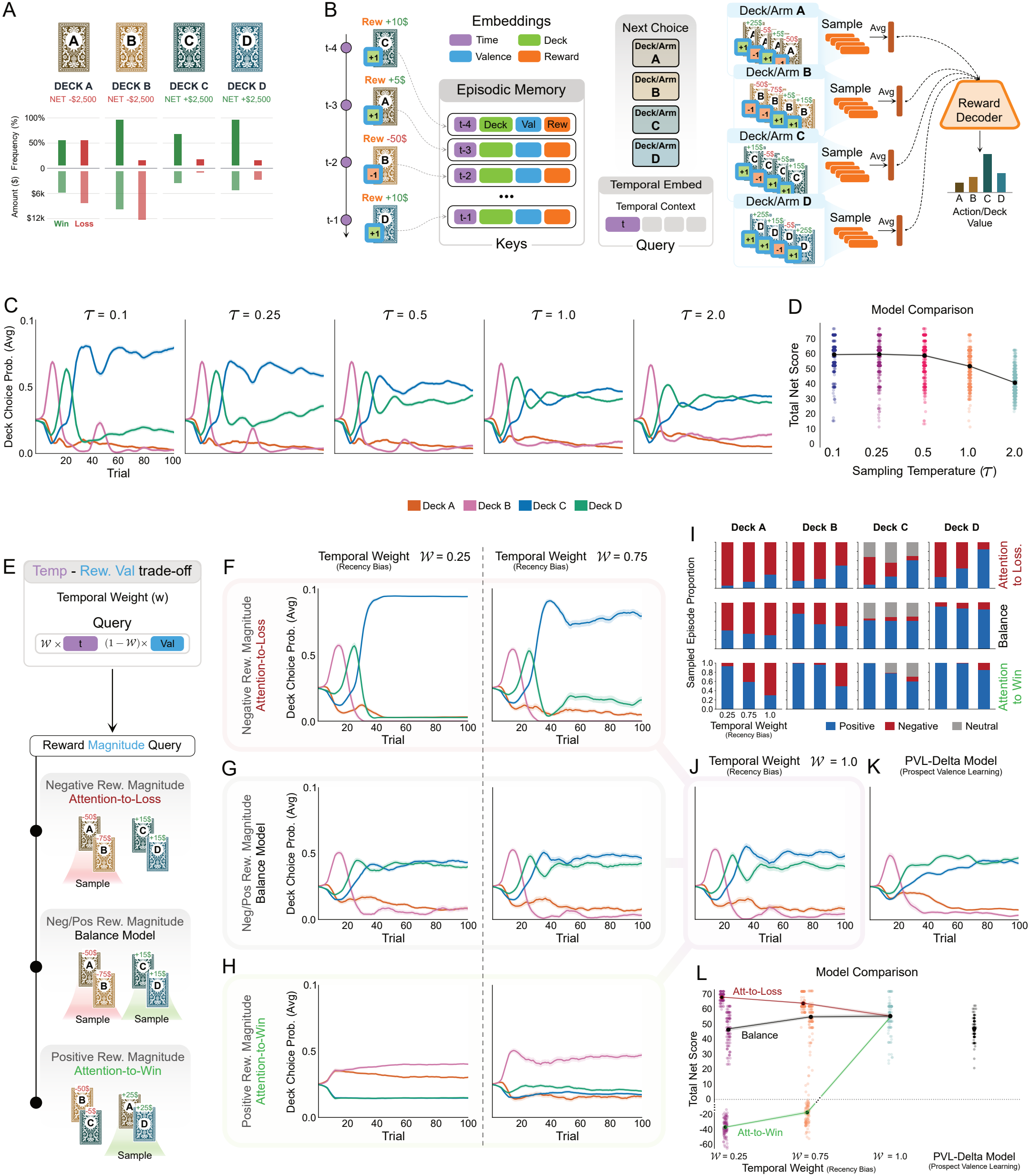
Recency and reward magnitude shape Iowa Gambling Task choices. **A**. Reward and punishment structure of the four decks. **B**. GENESIS-Value reconstructs deck values from *k* = 20 episodic samples per deck, drawn with replacement and decoded after averaging their reward embeddings. **C**. Deck choice probabilities for temporal-only retrieval across sampling temperatures. **D**. Corresponding Net Scores, defined as (*C* + *D*) − (*A* + *B*). **E**. Magnitude-selective queries combine temporal context with learned reward embeddings of the maximum punishment, maximum win, or both; balanced queries weight the two reward components equally. **F–H**. Deck choice probabilities across temporal weights under attention to loss (**F**), balanced attention to loss and win (**G**), and attention to win (**H**). **I**. Proportions of positive (blue), negative (red), and neutral zero-reward (grey) episodes sampled by deck and temporal weight for the three reward-magnitude models. Grey bars in Deck C reflect the presence of zero-reward outcomes in its payoff schedule. **J**. Deck choice probabilities with *w* = 1, isolating temporal retrieval. **K**. Choice probabilities of the PVL-Delta reference, with parameters *A* = 0.88, *w*_loss_ = 0.68, *a* = 0.25, and *c* = 1.27. **L**. Comparison of Net Scores across GENESIS retrieval conditions and the PVL-Delta reference.

In healthy participants, IGT performance generally shifts toward the advantageous decks (C and D), although substantial individual variability exists, with some participants favoring decks associated with infrequent punishments ^52,53^. Adaptive performance in healthy participants is reflected in positive IGT Net Scores, calculated as (C + D) − (A + B), which indicate a preference for decks that yield higher long-term returns. Persistent selection of disadvantageous decks offering large immediate rewards despite unfavorable long-term outcomes (A and B) has been documented in patients with ventromedial prefrontal cortex damage ^54^, highlighting the task’s relevance to the study of impaired decision-making ^55^. Beyond these findings concerning reward-sensitive decision-making, a separate line of evidence implicates episodic and declarative memory in the evaluation of past outcomes and the updating of deck preferences ^31,34^. Consistent with this contribution, individuals with amnesia and impaired declarative memory exhibit poorer IGT performance ^32,33^. We implemented the IGT task as a four-armed episodic bandit (Figure 6B). At each decision, one value per deck was reconstructed from that deck’s own episodic history using *k* = 20 replacement samples; the four reconstructed values were then passed to a softmax choice policy. Deck values were therefore constructed on the fly through episodic sampling, rather than represented by incrementally updated scalar estimates.

We tested the effects of different retrieval queries. We first restricted retrieval to temporal context and varied the choice temperature across five levels:

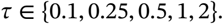

Five hundred agents were simulated at each temperature. Choice temperature was annealed from 5.0 to 0.25 over the first 10 trials and then held fixed, separating the retrieval-temperature manipulation from persistent changes in policy noise.

Across the task, for all levels of temperature, probability tended to move away from Deck A and, following an early period in which its large immediate gains made it attractive, also away from Deck B. The strongest effect of sampling temperature emerged in the balance between the two advantageous decks: low-temperature sampling produced a pronounced and stable concentration on Deck C, whereas high temperatures produced a broader distribution over the advantageous decks overall (Figure 6C). When considering the IGT Net Score, all models showed advantageous choice behavior, but low-temperature sampling models achieved higher scores, reflecting their early commitment to Deck C (Figure 6D).

We next tested a query that combines temporal information with reward sensitivity, whereby the probability of retrieving past episodes is modulated by the magnitude of their associated rewards and punishments. A strategy that prioritizes salient, reward-related episodes involving large-magnitude outcomes is consistent with the reward-sensitive choice patterns observed in the Iowa Gambling Task (IGT). We constructed reward-sensitive queries from the embeddings of the maximum win and maximum loss available in the task, corresponding to scaled outcomes *r*_max_ = +1.0 and *r*_min_ = −11.5, respectively, in combination with current temporal context (Figure 6E). We tested three reward-sensitive conditions: *attention to loss*, combining temporal context with the maximum-loss embedding; *attention to win*, combining temporal context with the maximum win embedding; and a *balanced* condition, combining temporal context with both embeddings. These queries probe similarity within a graded, learned reward representation.

For each memory retrieval, temporal and reward-sensitive cosine similarities were combined into a single retrieval score. Temporal context received weight *w* ∈ {0.25, 0.75, 1}, with complementary reward-sensitive weight 1 − *w*; in the balanced condition, this reward-sensitive contribution was divided equally between the win and loss components. All three model instances used *τ* = 0.25, *k* = 20, and 100 agents per temporal-weight cell. Choice temperature decreased from 5.0 to 0.5 over the first 10 trials and then remained fixed.

Attention-to-loss retrieval favored Deck C and suppressed Deck D most clearly when the reward-sensitive component carried greater weight in the query (Figure 6F). The balanced query instead supported an advantageous preference distributed across Decks C and D (Figure 6G). In contrast, attention-to-win retrieval sustained a strong attraction to decks offering larger immediate rewards, particularly Deck B, despite their unfavorable longrun returns (Figure 6H). This preference translated into negative Net Scores (Figure 6L), demonstrating that an excessive focus on highly rewarded experiences can produce persistently disadvantageous decisions. Notably, this behavior resembles the well-documented tendency to favor frequent immediate gains despite occasional large punishments, a characteristic pattern of IGT performance that can also emerge in healthy participants ^56^. Thus, altering the reward-sensitivity of the query was sufficient to shift GENESIS-Value from advantageous performance toward a strategy dominated by short-term reward – as an effect of retrieving different episodes (Figure 6I). Finally, when *w* = 1, reward sensitivity no longer contributed to retrieval, which was instead guided entirely by temporal context (Figure 6J), analogous to the temporal-context query described above.

As a reference, we also simulated a model widely used to account for performance in the Iowa Gambling Task: the Prospect Valence Learning model with delta updating (PVL-Delta), which relies on incremental learning rather than episodic sampling. We used a normative parameter combination that produces a preference for advantageous decks ^57^. The reference model favored Decks C and D collectively over A and B (Figure 6K), broadly resembling the balanced GENESIS-Value conditions (Figure 6G).

Finally, comparing Net Scores across models shows that the best-performing model is the attention-to-loss model at *w* = 0.25, where reward magnitude dominated temporal recency. The balanced-condition models achieved Net Scores comparable to that of the PVL-Delta reference model (Figure 6L).

In summary, these simulations show that GENESIS-Value supports adaptive decision-making in the Iowa Gambling Task (IGT) ^30^. Moreover, the same episodic sampling mechanism can give rise to qualitatively different reward-guided choice policies, depending on whether the retrieval query prioritizes temporal or reward-related dimensions. Preferential retrieval of large punishments produced the strongest Net Scores, whereas preferential retrieval of large wins led to disadvantageous choices dominated by immediate rewards. By contrast, jointly attending to wins and punishments yielded a more balanced preference for advantageous decks, resembling the behavior of the conventional PVL-Delta reference widely used to model the IGT ^57^.

## Discussion

A central question in understanding human decision-making is how people flexibly evaluate options and make choices across different contexts by drawing on their past experiences. In this study, we demonstrate that a single episodic model can support a variety of value-based decision-making tasks by enabling accurate value reconstruction and adaptive choice.

Classical approaches typically rely on stable value representations, which are compact but often need to be learned anew for different tasks. In contrast, GENESIS-Value constructs value on the fly by sampling and integrating information from past episodes, without relying on precomputed, stable value representations. Crucially, the model is based on a query–key–value mechanism that provides flexibility across decision-making contexts. Although the underlying episodic sampling process remains the same across tasks, the episodes retrieved and the way offers are evaluated depend on the query. This flexibility allows GENESIS-Value to address a range of scenarios examined in the literature: stimulus identity-based queries support the evaluation of experienced and novel stimuli ^14^; feature-based queries enable the estimation and choices between stimulus features ^17^; queries based on temporal context and reminder cues support sequential decision-making in bandit tasks ^18^; and queries combining temporal proximity with sensitivity to positive and negative outcomes support decision-making in the Iowa Gambling Task ^30^. Thus, the same episodic mechanism can flexibly guide memory retrieval and support different decision-making strategies depending on the query and the current decision context. This flexibility brings GENESIS-Value closer to accounts of human decision-making showing that people adapt their choices to different task demands and goals ^11–13^.

Some mechanisms in GENESIS-Value, such as sampling temperature, play a role analogous to the learning rate or recency-weight in models that learn stable value representations (e.g., through temporal-difference (TD) learning or averaging past rewards), as they influence the relative weighting of recent and more distant experiences. However, a key advantage of episodic retrieval in GENESIS-Value is that the temporal range of retrieved experiences can be adjusted dynamically at decision time. In contrast, TD learning relies on a learning rate that determines how strongly new experiences update existing value estimates, typically based on (implicit) assumptions about environmental volatility ^51^. If the environment proves more or less volatile than initially anticipated, the learning rate can be adjusted; however, in standard online TD learning, this primarily affects subsequent updates, because previous experiences have already been incorporated into the current estimates rather than retained as individual episodes. By preserving individual episodes, episodic systems allow past experiences to be re-evaluated under changing assumptions about environmental volatility. This enables agents to adapt to environments with different levels of predictability, which, as our simulations show, may benefit from different sampling temperatures. Along the same lines, GENESIS-Value provides a straightforward way to switch between decision strategies on the fly by changing the retrieval query – for example, shifting from prioritizing episodes associated with large wins to those involving large losses.

The theory advanced here does not necessarily contradict the existence or use of stable value representations in the brain, for which substantial supporting evidence exists ^1,5^(but see ^58^ for criticism). Rather, episodic retrieval may coexist with stable value representations, which could be relied upon, for example, in more predictable or stable contexts. Consistent with this possibility, Nicholas and Mattar ^17^ reported that participants relied on episodic memory to encode and retrieve specific experiences when they could not anticipate which features would be relevant for future decisions. In more stable contexts, by contrast, value representations may become consolidated or “cached” and used to guide choices. From this perspective, the present proposal fills an important gap by suggesting that episodic memory provides an additional source of flexibility when needed. Rather than replacing stable value representations, which have been a primary focus of neuroeconomics, episodic retrieval may complement them as an additional mechanism for guiding decisions.

GENESIS-Value builds on and extends influential accounts of episodic recall and decision-making in cognitive science and machine learning ^42,43^. It connects with classical models of memory recall and recognition, like SAM (Search of Associative Memory) ^59^, REM (Retrieving Effectively from Memory) ^60^, and the temporal context model ^61^, but uses a query–key–value mechanism which allows retrieval to be flexibly guided by different types of queries. Furthermore, GENESIS-Value shares with the framework of decision by sampling ^41^ the idea that choices can be guided by episodic sampling. However, decision by sampling focuses on a specific type of decision problem: comparing an offer with a reference value. Similarly, other episodic models have been developed to account for specific experiments, with limited ability to generalize to novel decision-making scenarios ^17,18^. As we have shown here, GENESIS-Value can support a broader range of decisions, by simply changing the query. Moreover, the model operates on naturalistic stimuli, such as images, and could potentially be extended to other modalities by incorporating suitable generative models. This makes it possible to test the frame-work directly in empirical settings that more closely resemble human decision-making tasks.

GENESIS-Value also connects to machine learning approaches that make decisions based on individual instances or memories rather than relying purely on generalized rules or abstract models. Classical examples in machine learning are case based reasoning ^62^ and instance-based or memory-based learning ^63^. Another way episodic memory has been incorporated into machine learning methods is through experience replay, in which stored experiences are repeatedly sampled from a buffer, loosely inspired by hippocampal replay in biological organisms ^64–67^. Experience replay can improve the sample efficiency of RL methods ^68–72^, mitigate catastrophic forgetting in continual learning ^73^, and provide approximate representations of value functions ^44,74^. Other approaches use episodic memory to generate intrinsic rewards for exploration ^75^, construct cognitive maps for context-aware action selection ^76^, or improve the performance and sample efficiency of neural networks in tasks involving memory and decision-making ^77^. However, these approaches do not use a flexible query–key–value mechanism to guide episodic retrieval for value-based decision-making. Although some episodic methods support learned and flexible memory retrieval, they primarily target improved world-model predictions rather than value-based choice ^78^. More broadly, these approaches typically optimize specific objectives, such as reward maximization, sample efficiency, or prediction accuracy, while their relationship to empirical findings on the role of episodic memory in human value-based decision-making remains unclear.

From a neurophysiological perspective, the episodic frame-work presented here is consistent with a large body of research highlighting the role of episodic memory in flexibly constructing and evaluating possible future events and situations ^79–83^. In GENESIS-Value, value construction is an integrative process involving several components: semantic processing (stimulus encoding and decoding, and semantic embeddings), potentially associated with cortical regions; episodic processing (memory encoding and retrieval), potentially associated with the medial temporal lobe; and valuation (reward encoding and decoding, and reward embeddings), potentially involving structures such as the orbitofrontal cortex and ventral striatum. This integrative perspective is consistent with classical accounts of complementary memory systems in the cortex and hippocampus ^84–86^, as well as more recent accounts that revisit these ideas within a generative modeling framework ^49,67,87–89^. Relatedly, human neuroimaging studies reported the involvement of both the hippocampus and the medial prefrontal cortex during the construction and evaluation of novel choice offers ^14^, the synchronization of medial temporal lobe and prefrontal rhythms during decision making ^90^, and the contribution of hippocampus-striatal connections to the valuation of novel stimuli ^91^. However, the extent to which the mechanisms implemented in GENESIS-Value correspond to neural dynamics, and which relevant processes are not captured by the current framework, remain open questions.

The retrieval-augmented generation (RAG) mechanism used by GENESIS-Value, which originates in machine learning ^48^, is compatible with evidence that the brain may employ key–value mechanisms to organize and retrieve information from memory ^50^. This assumption is shared by other models of episodic memory construction involving images ^49^ and stories ^92^. However, how the brain implements biological mechanisms analogous to RAG remains an open question.

The model presented here has several limitations that future research should address. First, although GENESIS-Value can flexibly guide episodic retrieval using different queries, it does not specify how or under what conditions a particular query is selected (beyond cases in which the query is determined by the experimental task). Future work could use the framework to investigate how people select or learn contextually appropriate queries, how query selection differs across decision contexts – for example, when evaluating a single offer versus choosing among multiple offers – and how it varies across individuals. It could also examine whether human query selection is optimal and under what circumstances.

Second, the model does not specify how the sampling budget or decision criterion is determined. These processes could be potentially incorporated using established approaches to value-based decision-making, such as evidence accumulation to a decision threshold ^93^, or by adjusting the sampling budget according to the expected value of information or other cost–benefit considerations ^45–47^.

Relatedly, the model presented here makes no specific assumptions about which episodes are encoded into memory during decision-making or the format in which they are stored. Rather, it requires only that memory contains sufficient information to guide retrieval and value construction in ways that support the decision at hand. Previous studies have shown that the encoding of episodes depends on their salience and occurs preferentially at meaningful event boundaries ^94^. Future studies could incorporate these findings to develop more biologically plausible mechanisms of episodic encoding, in which the selection and storage of experiences depend on their salience and the demands of the ongoing task.

Finally, our primary goal was to demonstrate that a single episodic model can address a range of scenarios used to study episodic contributions to human value-based decision-making ^14,17,18^, as well as other tasks involving flexible choice, such as the Iowa Gambling Task ^30^. Although our results qualitatively reproduce behavioral signatures associated with human episodic decision-making, we do not directly compare the model’s predictions with human behavioral data. Future work should therefore evaluate the model against empirical data to assess how well it captures human behavior, identify the conditions under which its predictions align with or diverge from observed choices, determine which parameter ranges (e.g., sampling budgets and temperatures) best reproduce human behavior, and further constrain the mechanisms underlying episodic value-based decisions. By operating on naturalistic stimuli, such as images, the model can be evaluated using materials comparable to those employed in human experiments, facilitating direct comparisons with empirical findings.

In summary, GENESIS-Value provides a unified framework for episodic value-based decision-making and a potential complement to theories emphasizing stable value representations. By demonstrating how a common retrieval mechanism can support diverse forms of valuation and choice, the framework offers a possible account of the flexibility of human decision-making.

## Methods

### GENESIS-Value architecture and episodic retrieval

GENESIS-Value extends the model of episodic and semantic interactions in memory encoding and retrieval of D’Alessandro et al. ^49^ to support value-based decision-making. In the simulations reported here, the cortical representational models were trained before the behavioral experiments and then frozen. Episodic memory was implemented as an explicit retrieval-augmented store of compressed events ^48^ rather than as an incrementally updated action-value table. This design isolates the consequences of changing what information is used to access memory, how selectively memories are sampled, and how many retrieved episodes are recombined at decision time.

#### Cortical stimulus and reward representations

The stimulus model was a conditional variational autoencoder (CVAE) trained on a restricted Shapes3D image set. Each 64 × 64 RGB image depicted an object described by four categorical factors: *object shape* (four classes: cube, sphere, cylinder, and pill), *object color* (five classes), *floor color* (five classes), and *wall color* (five classes), yielding 4 × 5^3^ = 500 possible combinations of semantic features. The dataset was divided into an 80/20 train–test split stratified by the joint combination of these factors (random seed 44). Unless otherwise specified, all Shapes3D-based simulations used the same pretrained CVAE, with task-specific restrictions on the stimuli when required. The CVAE weights remained frozen throughout the behavioral simulations.

The CVAE separates image information into a continuous latent representation and four learned, factor-specific conditioning embeddings. Its encoder approximates a Gaussian distribution *q*_*ϕ*_(*z* ∣ *x*) over the image latent *z*, whereas the decoder reconstructs the image using both *z* and the four conditioning embeddings. Importantly, these embeddings are learned from the categorical factor labels during training; they are not assumed to emerge automatically as disentangled dimensions of the image latent. Conditioning the decoder on each factor separately makes object shape and the three color features individually available for later episodic queries. The continuous latent retains additional visual information needed to reconstruct the particular image.

For image *x*_*t*_, we use the posterior mean *z*_*t*_ = *µ*_*ϕ*_(*x*_*t*_) at inference, together with the four learned factor embeddings:

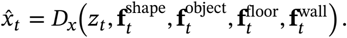

Here, **f**^shape^ explicitly represents *object shape*, **f**^object^ represents *object color* (not object identity), and **f**^floor^ and **f**^wall^ represent floor and wall color, respectively. The conditioning embeddings modulate the decoder through feature-wise linear modulation (FiLM). During training, image reconstruction loss was combined with a Kullback–Leibler penalty that regularized the Gaussian latent distribution. The CVAE used three spatial down-sampling stages, four latent channels, a base width of 32 channels, four conditioning channels per factor, and KL weight 0.025. It was trained for 30 epochs with batch size 32 and learning rate 1.5 × 10^−3^. The trained CVAE was then frozen. Episodic simulations stored the posterior-mean image latent and the four factor embeddings separately, permitting queries based on any combination of these representations. In the reminder-bandit task, the flattened image latent and four factor embeddings were also concatenated to form a single stimulus-retrieval cue.

Scalar rewards were represented by a separately trained encoder–decoder network. The encoder consisted of a scalar-to-4-unit linear projection followed by a ReLU and an 8-dimensional latent projection; the decoder was a single affine map from the 8-dimensional reward embedding to a scalar. The model was optimized with mean-squared reconstruction loss for 50 epochs using AdamW, batch size 8, and learning rate 10^−2^, and was frozen during all behavioral simulations. Separate checkpoints with the same architecture were trained for the standard reward range used by the Shapes3D experiments and for the scaled IGT payoff range. For an experienced outcome *r*_*t*_,

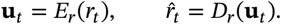

Because *D*_*r*_ is affine, decoding the mean of a set of reward embeddings is equivalent to averaging their individually decoded rewards. Thus, any discrepancy between the decoded aggregate and the corresponding mean scalar reward reflects reward-autoencoder reconstruction error rather than the averaging operation itself.

#### Temporal context and episode formation

Each stored event was assigned a 64-dimensional unit-norm temporal-context vector. Temporal context followed a stochastic trajectory with a fixed angular increment *γ* = *π*/100:

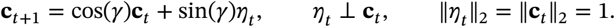

At every step, *η*_*t*_ was generated as a random unit vector in the subspace orthogonal to the current context. Successive contexts therefore changed smoothly while remaining on the unit hypersphere, providing a temporally ordered retrieval key without modifying the reward stored in an episode.

In the feature-rich experiments, an episode was represented as

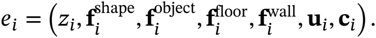

Here, *z*_*i*_ is the continuous image latent; 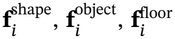, and 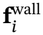 are the embeddings for object shape, object color, floor color, and wall color; **u**_*i*_ is the encoded reward; and **c**_*i*_ is the temporal context at encoding. Sequential tasks additionally associated the episode with the selected arm or deck. Action identity was used to define action-specific candidate pools rather than as a similarity term in the retrieval score. Thus, retrieval reconstructed the value of an action only from episodes in which that action had previously been selected.

#### Query–key matching and stochastic retrieval

A decision-time query *q*_*t*_ could contain any subset of the stored key components, including a semantic feature, current temporal context, a reminder stimulus, or an experienced reward magnitude. For each active component *m*, query–key similarity was cosine similarity,

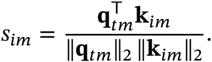

When multiple components were combined, they were first placed on a common candidate-relative scale. Feature-only and temporal–reward trade-off simulations standardized each active similarity component across the current candidate set,

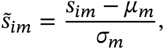

with a zero vector used when *σ*_*m*_ = 0. The reminder-bandit used component-wise rank normalization: candidates were ranked from highest to lowest similarity separately for each component and mapped linearly to equally spaced scores in approximately [−1, 1]. In single-component temporal-only conditions, the resulting score vector was standardized across candidates before sampling. These normalizations prevent differences in the numerical scale of heterogeneous similarity components from implicitly changing their intended query weights.

For the active component set *A*, normalized similarities were combined as

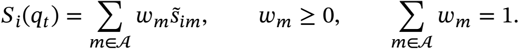

A zero-weight component was excluded from the score. Candidate accessibility was then converted to a sampling distribution by

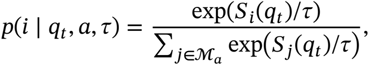

where *ℳ*_*a*_ is the candidate set for action *a*, or the complete session memory in non-sequential feature experiments. Low *τ* increases the influence of small score differences and concentrates retrieval on the most accessible events; high *τ* approaches a flatter distribution. Unless noted otherwise, episodes were sampled with replacement, allowing a highly accessible episode to contribute multiple times to the same reconstructed value.

For a sampled index multiset *I*_*a*_, reward embeddings were averaged and decoded,

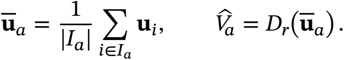

The sampling budget therefore controls the Monte Carlo precision of the reconstructed reward aggregate, whereas *τ* controls the accessibility distribution from which that aggregate is drawn. In the two-armed bandit, the number of draws from an arm during early trials was capped by the number of episodes available for that arm, |*I*_*a*_| = min(*k*, |*ℳ*_*a*_|), while sampling remained with replacement. In the IGT, exactly *k* samples were drawn from every nonempty deck memory; unseen decks were assigned an initial reconstructed value of zero.

For the two-armed sequential tasks, reconstructed values were mapped to choice through

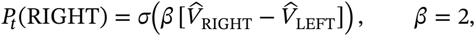

with stochastic Bernoulli choice. For the IGT, a four-way softmax policy was used; its trial-dependent temperature is specified below for each condition. After a choice, the observed outcome and current stimulus/context representations were encoded and appended to episodic memory, completing the retrieval–choice– encoding loop.

Across all simulations, queries were matched against episodic keys, while the corresponding values provided the embeddings used for task-specific computations. In this implementation, for simplicity, keys and values share the same stimulus and reward representations, differing only in the temporal-context embedding included in the keys but not in the values (see ^49^ for an alternative formulation in which keys are more compressed compared to values).

### GENESIS-Value instantiations across experiments

#### Value reconstruction for seen and unseen stimuli

To examine generalization from episodic experience, we created 125 independently seeded sessions of cube images varying in object, floor, and wall color. In each session, one image was selected for every combination of five levels of these three factors (5^3^ = 125 combinations). Each color level contributed a value from {−2, −1, 0, 1, 2}, and the ground-truth reward of stimulus *s* was the mean of its three color contributions,

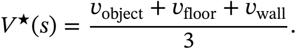

A target conjunction was selected within each session. In the *seen* condition, all 125 stimuli and their rewards were stored in episodic memory; in the matched *unseen* condition, the target and its reward were withheld, leaving 124 episodes. The query always represented the same target stimulus but never included its reward. It weighted the image latent, object-color embedding, floor-color embedding, and wall-color embedding equally (0.25 each); object shape and temporal context had zero weight. Component similarities were standardized across candidate episodes, and retrieval used *τ* = 0.25 with replacement. Sampling budgets *k* ∈ {1, 3, 5, 10, 20} were evaluated with 500 independent retrieval repetitions per session and condition. The sampled reward embeddings were averaged and decoded on each repetition. Figure 2 presents reconstruction at *k* = 20, comparing decoded values with ground-truth target rewards across matched seen and unseen sessions.

#### Feature-conditioned reward reconstruction

The feature-conditioned experiment isolated semantic control of reward reconstruction. Twenty independently seeded sessions were sampled from the held-out Shape3D test set, with 200 unique images per session. Object shape was exactly balanced across its four levels, and object, floor, and wall colors were balanced within shape across their five levels. Rewards {−2, −1, +1, +2} occurred equally often, so the complete-session reward mean was zero. Reward permutations were accepted only if every factor level had a nonzero reward sum, the complete set of factor-level reward sums contained both positive and negative values, and the mean reward of cube episodes was at least 0.5. The accepted sessions produced the cube-conditioned reward distribution reported in Section .

Each session was encoded once and stored as factorized episodes. The query contained the learned object-shape embedding for cube and a terminal temporal context, but temporal weight was set to zero; all other feature components also had zero weight. All 200 episodes remained eligible for sampling, so semantic selectivity arose from the query–key distribution rather than from hard filtering of the store. Object-shape similarity was standardized over the session candidates, sampling temperature was fixed at *τ* = 0.25, and *k* ∈ {1, 3, 5, 10, 20} replacement draws were tested. Each session-by-budget condition was repeated 500 times using independent retrieval seeds.

For each repetition, the sampled reward embeddings were averaged and decoded to obtain *r*_cube_. Session-level performance was summarized after averaging the 500 reconstructed values. The normalized reconstruction score plotted in Figure 3C was

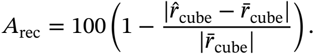

Across-session means and standard errors for Figure 3C were estimated by 5,000 bootstrap resamples of the independent session-level summaries (random seed 44).

#### Flexible feature comparison

The feature-comparison experiment used the same 200-item balanced session construction but required two independent semantic queries. Forty sessions were generated in two equal cohorts whose oracle choice was either cube or sphere. Baseline rewards again contained equal counts of −2, −1, +1, and +2. For each cohort, the target absolute cube– sphere mean-reward gap was evenly spaced from 0.1 to 1.0. After a baseline reward permutation, an equal and opposite offset was added to cube and sphere outcomes so that the realized signed gap exactly matched the scheduled value,

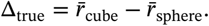

Assignments were retained only when all factor-level reward sums were nonzero and the sign of Δ_true_ matched the required oracle feature.

Cube and sphere were queried separately using their learned object-shape embeddings. Temporal weight was zero, all session episodes remained eligible, component similarities were standardized across candidates, *τ* = 0.25, and the same five sampling budgets were used. Five hundred paired retrievals were performed per session and budget. A decision selected cube when 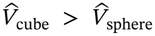, sphere when the inequality was reversed, and returned a tie for exactly equal reconstructed values; ties were scored as incorrect. Choice accuracy was the fraction of the 500 paired decisions matching the session’s oracle feature. Error bars in the budget-level summary were obtained by 5,000 bootstrap resamples over independent sessions (random seed 44).

#### Temporal selectivity in the volatile two-armed bandit

The sequential task followed the episodic-sampling logic of Bornstein et al. ^18^ while using GENESIS-Value representations. Each environment contained 130 choice trials. The left-arm reward probability began at either 0.6 or 0.4 with equal probability and then followed a Gaussian random walk with innovation standard deviation 0.025, reflected at the bounds [0.3, 0.7]. The right-arm probability was constrained to 1 − *p*_left,*t*_. Reward outcomes were binary, *r*_*t*_ ∈ {−1, +1}. A trial-unique Shape3D semantic class was assigned to each choice trial, and the event stored after choice contained the chosen arm, its outcome, the encoded stimulus, and temporal context.

The first 20 trials were random-choice warm-up trials and were nevertheless stored in episodic memory. On trials 21–130, the current temporal vector was the sole active retrieval component. For each arm, only episodes generated by previous choices of that arm entered its candidate set. The sampling budget was *k* = 20, with the early-memory cap described above, and *τ* ∈ {0.1, 0.25, 0.5, 1, 2}. One hundred independently generated environments were crossed with 50 independently seeded agents per environment and temperature condition. The same environment schedule was therefore shared across the temperature comparison, while retrieval, choices, and realized binary outcomes remained stochastic at the agent level.

Environmental predictability was computed solely from the reward-probability schedule. Let

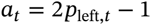

denote the signed left-arm advantage. For an early window of length *N*, we defined

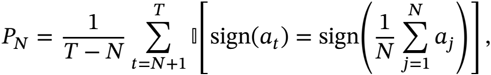

excluding exact ties. The environment score was the mean of *P*_*N*_ over *N* ∈ {20, 30, 40}. It therefore measures the fraction of later trials for which the early direction of reward advantage remains valid. For Figure 4H, environments were divided into equal-sized low-, intermediate-, and high-predictability groups by tertiles of this score. Across-environment associations in Figure 4F were summarized with Pearson correlations; their 95% intervals were obtained from 5,000 environment-level bootstrap resamples.

#### Reminder-biased episodic valuation

The reminder experiment retained the same bandit dynamics, temporal contexts, actionvalue reconstruction, and logistic policy but added stimulus information to selected retrieval queries. The schedule contained 24 reminder trials per 130-choice environment. For every trial-unique Shape3D semantic class, two distinct images were reserved: one was shown and stored on the original choice trial, whereas the other was held out for possible later use as a reminder. Reminder trials were sampled without replacement from eligible later trials. Each reminder targeted a distinct earlier choice 20–70 trials in the past, and the reminder image came from exactly the same four-factor semantic class as the target while having a different HDF5 image identity. Consequently, a reminder matched the target semantically without repeating the original visual sample.

On non-reminder trials, retrieval was temporal-only. On reminder trials, the query contained both current temporal context and the encoded held-out reminder stimulus. The reminder-bandit placed temporal and stimulus similarities on a common component-wise rank scale before combining them,

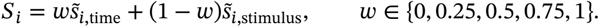

Thus, *w* = 0 gave the reminder complete control over query–key matching and *w* = 1 reproduced the temporal-only retrieval rule. The sampling budget was fixed at *k* = 20 with replacement and *τ* ∈ {0.1, 0.25, 0.5, 1, 2}. Fifty independent environments with 50 agents per environment were simulated.

For a target episode with sampling probability *p*_target_ among *N*_*a*_ eligible memories for its arm, accessibility enrichment was defined relative to uniform sampling as

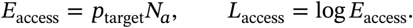

Positive *L*_access_ indicates that the reminder target was more accessible than under a uniform distribution over the same action-specific memory pool. To quantify propagation from retrieval into valuation, let *s*_*a*_ = +1 for a right-arm target and −1 for a left-arm target. Signed value alignment was

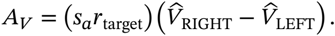

Positive values therefore correspond to a reconstructed value difference that favors repeating the target action after a win or switching away from it after a loss. The choice-level endpoint was whether the post-reminder action repeated the target action, summarized separately for target wins and losses relative to a 0.5 reference probability. Confidence intervals in Figure 5 were generated by 5,000 bootstrap resamples over independent environments (random seed 44), preserving agents and reminder trials within their generating environment.

#### Iowa Gambling Task: temporal and reward-magnitude retrieval

We modeled the Iowa Gambling Task (IGT) as a four-armed episodic bandit using a fixed 100-row payoff schedule for Decks A–D (Figure 6A). Decks A and B provide larger immediate gains but disadvantageous long-run returns; Decks C and D provide smaller immediate gains but advantageous returns. Each deck had its own draw pointer: choosing deck *d* revealed its next unused payoff and advanced only that deck’s pointer. Payoffs were multiplied by 0.01 before reward encoding. Each agent made 100 choices. On every trial, GENESIS-Value separately queried the previous episodes associated with each deck, sampled *k* = 20 episodes with replacement from each nonempty deck memory, averaged their reward embeddings, and decoded one value 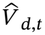 per deck. Unseen decks were assigned an initial value of zero.

Choices were then sampled using the four-way softmax

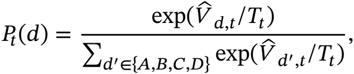

where *T*_*t*_ denotes *choice* temperature, distinct from the episodic *retrieval* temperature *τ*. The experienced reward and temporal context were stored after each choice. No fixed deck-value estimates were learned or maintained by GENESIS-Value.

We examined two related retrieval manipulations using this common task and memory mechanism. First, in the *temporal-only* condition, the current temporal context was the only query component. We varied sampling temperature over *τ* ∈ {0.1, 0.25, 0.5, 1, 2} and simulated 500 independently seeded agents per temperature. The choice temperature decreased linearly from 5.0 on trial 1 to 0.25 on trial 10 and then remained fixed. This experiment isolates the influence of retrieval recency on deck preference (Figure 6C,D).

Second, we tested three *reward-magnitude-selective* queries that combined current temporal context with learned reward embeddings. Instead of attaching a categorical win/loss label to each episode, we used the same continuous reward representations already stored for reconstructing payoff values. We encoded the most negative and most positive scaled outcomes present in the IGT schedule, *r*_−_ = −11.5 and *r*_+_ = +1.0, to obtain the two query cues **u**_−_ = *E*_*r*_(*r*_−_) and **u**_+_ = *E*_*r*_(*r*_+_). For each stored episode *i*, we computed cosine similarities to current time (*s*_*i*,time_), the maximum-punishment cue (*s*_*i*,−_), and the maximum-win cue (*s*_*i*,+_). Each active similarity component was standardized separately across episodes available for the queried deck. With temporal weight *w*, the three retrieval models used

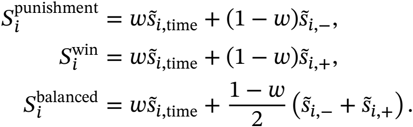

Thus, the *attention-to-loss* model preferentially accesses outcomes similar to the largest loss, the *attention-to-win* model preferentially accesses outcomes similar to the largest gain, and the *balanced* model weights both magnitude cues equally (Figure 6E–H). These are graded similarities in the learned reward space, not binary restrictions on which episodes can be retrieved: outcomes of any sign remain eligible. In all three models, *w* ∈ {0.25, 0.75, 1}, sampling temperature was fixed at *τ* = 0.25, and 100 agents were simulated per temporal-weight condition. The choice temperature decreased from 5.0 to 0.5 during the first 10 trials and remained at 0.5 thereafter. At *w* = 1, reward magnitude has zero query weight, providing a temporal-only comparison *within* the unified magnitude-selective experiment (Figure 6J). The separately simulated temporal-only experiment above used its original query normalization and a different final choice temperature.

The principal behavioral measure was the conventional IGT Net Score, calculated over the complete task as

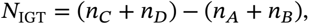

where *n*_*d*_ counts choices from deck *d* in the relevant period. Deck choice probabilities, Net Scores, and retrieval-outcome proportions were aggregated across agents. Uncertainty estimates were obtained using 10,000 agent-level bootstrap resamples with percentile 95% intervals (random seed 44).

For comparison, we simulated a conventional Prospect Valence Learning model with Delta updating (PVL-Delta) ^57^. Unlike GENESIS-Value, PVL-Delta maintains an incrementally updated expectancy *Q*_*d,t*_ for each deck. After choosing deck *d* and receiving scaled net payoff *r*_*t*_, its prospect utility and chosen-deck update are

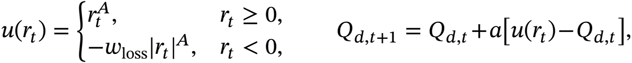

while the other deck expectancies remain unchanged. Here *A* controls outcome sensitivity, *w*_loss_ controls the relative influence of losses, and *a* is the learning rate. Deck choices follow *P*_*t*_(*d*) *∝* exp(*θ*_*t*_*Q*_*d,t*_), with base response sensitivity *θ* = 3^*c*^ − 1 determined by the consistency parameter *c*. We used the fixed values *A* = 0.88, *w*_loss_ = 0.68, *a* = 0.25, and *c* = 1.27, drawn from the advantageous-choice example of Steingroever et al. ^57^ , rather than fitting parameters to GENESIS-Value or human data. All initial deck expectancies were zero. To match the GENESIS magnitude-model exploration schedule, response sensitivity was reduced during the first 10 trials by *θ*_*t*_ = *θ* (0.5/*T*_*t*_), where *T*_*t*_ decreases from 5.0 to 0.5; thereafter *θ*_*t*_ = *θ*. One hundred independent PVL-Delta agents experienced the same four-deck schedule and payoff scaling as the magnitude-selective models. Its deck-choice probabilities and Net Scores were analyzed alongside the GENESIS-Value conditions (Figure 6K,L).

## Acknowledgements

This research received funding from the European Research Council under the Grant Agreement No. 820213 (ThinkAhead), the European Union under the Horizon Europe research and innovation programme (Grant Agreement No. 101298363, Project OPERA), and the Ministry of University and Research, PRIN 20229Z7M8N. The funders had no role in study design, data collection and analysis, decision to publish, or preparation of the manuscript. Views and opinions expressed are however those of the author(s) only and do not necessarily reflect those of the European Union or European Health and Digital Executive Agency (HaDEA). Neither the European Union nor the granting authority can be held responsible for them. We used a Generative AI model to correct typographical errors and edit language for clarity.

